# Archaeal type IV pili stabilize *Haloferax volcanii* biofilms in flow

**DOI:** 10.1101/2023.01.20.524888

**Authors:** Pascal D. Odermatt, Phillip Nussbaum, Sourabh Monnappa, Lorenzo Talà, Zhengqun Li, Shamphavi Sivabalasarma, Sonja-Verena Albers, Alex Persat

## Abstract

Biofilms represent a prevalent lifestyle of unicellular organism that confers protection to external challenges. The mechanisms by which archaea form biofilms are however not entirely clear. *H. volcanii* is an extremely halophilic euryarchaeon that commonly colonizes salt crust surfaces. *H. volcanii* produces long and thin appendages called type IV pili that are known to play a function in surface attachment and biofilm formation in archaea and bacteria. Here, we used biophysical experiments to identify critical function of type IV pili in the mechanical integrity of *H. volcanii* biofilms. Using interferometric scattering microscopy (iSCAT) to non-invasively visualize T4P in live cells, we find that piliation varies across mutants expressing single pilin isoforms. Using microfluidic experiments, we found that the adhesive strength of these mutants correlates with their extent of piliation. We found that in flow, *H. volcanii* forms clonal biofilms that extend in three dimensions. Expression of PilA2, a single pilin isoform, is sufficient to maintain normal levels of piliation and form biofilms with a structure indistinguishable from WT. Furthermore, we found that fluid flow is a crucial determinant of biofilm integrity: in the absence of flow, biofilms lose cohesion and tend to disperse in a density-dependent manner. Overall, our results demonstrate that T4P-surface and possibly T4P-T4P interactions promote biofilm formation and integrity, and that flow is a crucial ingredient regulating archaeal biofilm formation.

## Introduction

In their natural environments, many archaea experience extreme physicochemical selective pressures including high salinity, low pH or elevated temperatures^1,2^. Archaeal species have evolved to grow in and adapt to these harsh selective pressures. Multicellular communities of contiguous cells attached to solid surfaces called biofilms represent a common bacterial strategy to improve resilience in adverse environments^3–5^. Sheltering from external environmental stressors improves the fitness of biofilm-dwelling cells, making biofilm the most prevalent lifestyle for terrestrial microbes. While bacterial biofilms have been under intensive investigation due to their importance for human health and industry, whether archaeal biofilms follow similar assembly rules remains unknown^6,7^.

Biofilms typically form as single cells attach on a surface and divide while maintaining cohesion with adjacent cells, thereby forming discrete multicellular aggregates. To maintain biofilm cohesion, single cells secrete extracellular polymeric substances (EPS) that connect contiguous cells and the entire community. Investigations of bacterial model systems such as the pathogens *Pseudomonas aeruginosa* and *Vibrio cholerae* have revealed their basic principles of biofilm morphogenesis and the associated emergent properties^8^. These two species abundantly secrete polysaccharides that form a gel matrix cementing cells together. Other species including *Neisseria meningitidis* and *Escherichia coli* use proteinaceous filaments such as pili and fimbriae to maintain a cohesive multicellular structure^9^. In archaea, various *Sulfolobus* species, a variety of halophilic strains and methanogens can form surface-associated communities reminiscent of biofilms^3,10–13^. However, the basic principles of archaeal biofilm formation and robustness to external forces are unknown^3,14^.

In addition to cell-cell cohesion, initial surface attachment is a critical to the initiation and stability of biofilms. Direct cell body adhesion promotes attachment, but is very sensitive to surface physicochemical properties^15^. Many bacteria solve this by employing filaments that can reach and anchor to the surface at a distance. Type IV pili (T4P) are micrometer-long protein filaments extending from the cell surface that many times play critical roles in early bacterial biofilm formation. *P. aeruginosa* uses T4P to attach to surfaces, and their successive extension retraction cycles to explore the surface which ultimately induces aggregation^16,17^. In a similar manner, *V. cholerae* attaches to surfaces with T4P to form biofilms^8,18^. In *Neisseria meningitidis*, T4P promote both cell-surface and cell-cell attachment^19^. Archaea exhibit a wide variety of T4P filaments which have been shown to be important for surface adhesion, cell-cell recognition, sugar uptake and motility^11,20,21^. For example, *Sulfolobus acidocaldarius* exhibits two different T4P, the archaeal adhesive pilus^22^ (aap) and the UV inducible pilus^23^ (Ups), which both mediate static biofilm formation and affect their morphology^22^. The archaellum mainly plays a role in the dispersal of the *S. acidocaldarius* biofilm. *Haloferax volcanii* also produces T4P, which promote adhesion to the surface of abiotic materials^24,25^.

Archaeal and bacterial T4P machineries are evolutionary related and follow the same assembly steps^26^. Interestingly, *H. volcanii* expresses not one but six pilin isoforms, PilA1 to PilA6 encoded in four separate operons (**Fig. 1A**). The contributions of each of these isoforms to T4P formation and function are unclear. A mutant lacking all six isoforms cannot attach to surfaces, while strains expressing any of the single isoforms recovers some ability to attach, suggesting they are able to assemble T4P^25^. These observations indicate that *H. volcanii* uses T4P in initial surface attachment, thereby potential playing a role in biofilm formation.

**Figure 1:**
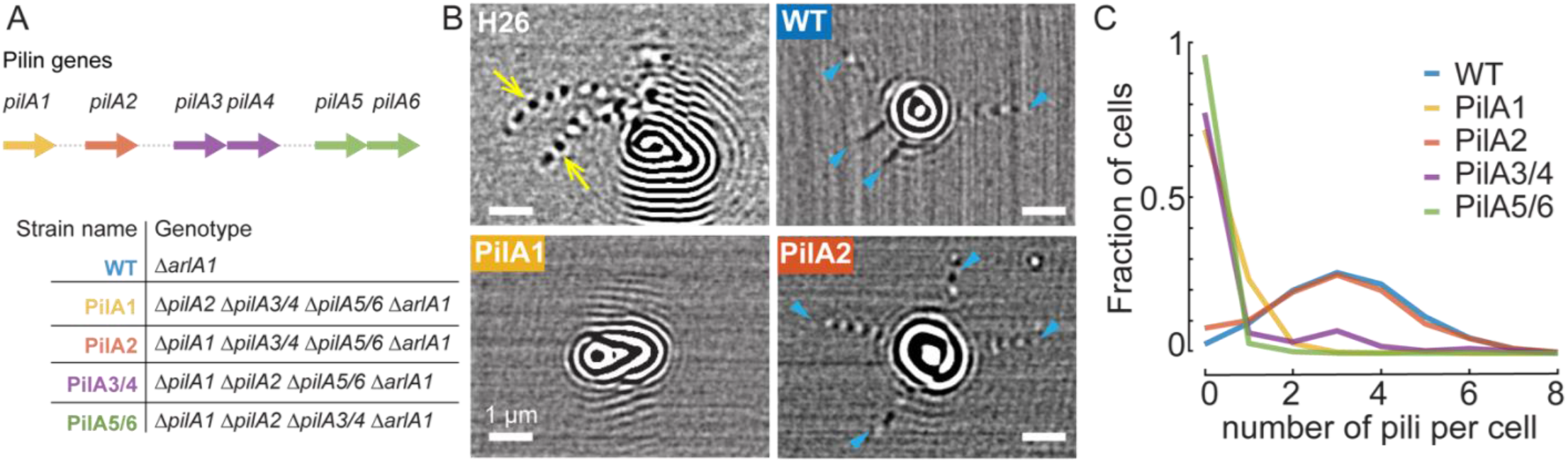
The PilA2 pilin subunit isoform is sufficient to produce type IV pili in *H. volcanii*. **A)** Description of strains expressing single pilin isoforms. **(B)** Representative iSCAT images of *H. volcanii*. (Top left) Archaella are visible in H26 (yellow arrows). (Top right) WT (archaellum-less H26 *ΔarlA1*) expressing all six PilA isoforms displays multiple T4P (blue arrowheads). (Bottom left) PilA1 typically does not produce T4P. (Bottom right) T4P are commonly visualized at the surface of PilA2. **(C)** Distribution of T4P numbers per cell for strains expressing different PilA subunit isoforms. The median T4P number for WT and PilA2 is 3, while all other strains have zero median T4P number.

Here, we rigorously investigate the functions of *H. volcanii* T4P during biofilm formation under flow. We first resolve the contributions of each pilin isoform in T4P biogenesis and surface attachment. Then, we focus on *H. volcanii* biofilm morphogenesis and show that flow promotes the cohesion of the multicellular community, suggesting a role of T4P in maintaining cell-cell cohesion in a force-dependent manner.

## Results

### The PilA2 pilin isoform is sufficient to maintain WT piliation

To visualize T4P at the surface of live *H. volcanii*, we used interferometric scattering microscopy (iSCAT) microscopy. We had previously employed iSCAT for the dynamic visualization of T4P in live *P. aeruginosa* cells at high temporal resolution^27^. Here, the sensitivity of iSCAT allowed us to resolve these filaments in live archaea without the need of labels. In iSCAT images, the H26 WT strain displayed many archaella which scatter light strongly, obstructing visualization of T4P (**Fig. 1B, supplementary movie 1**). To reveal T4P, we therefore imaged cells in a background lacking archaella by deletion of the *arlA1* gene. We subsequently refer to this background as WT. To test the contributions of each pilin isoform in T4P biogenesis, we generated mutants with a single pilin type by deleting three out of the four pilin operons among *pilA1, pilA2, pilA3/4* and *pilA5/6* (**Fig 1A**). We refer to a mutant strain with all pilins deleted but pilin *n* as PilA*n*.

On iSCAT images, T4P appear as long and thin filaments emanating from the strongly-scattering cell body (**Fig. 1B**). WT had many T4P, while mutants differed in their distributions of T4P number per cell. We found that more than 90% of WT and PilA2 had T4P, both with a median of 3 per cell (**Fig. 1C**). By contrast, more than 60% of PilA1, PilA3/4 and PilA5/6 cells did not visibly produce T4P under iSCAT imaging. This demonstrates that the expression of the pilin subunit PilA2 alone is sufficient to maintain WT-level of piliation. In contrast, expression of any other pilin subunit alone severely limits piliation, but does not abolish the ability to form T4P. We noticed that most T4P attach to the surface by their tip during the entire visualization time (typically 10 s). Out of the >1,500 T4P we visualized, we could not find any obvious evidence for retraction. This suggests that T4P may not power surface-associated motility in *H. volcanii*, consistent with the fact that archaeal genomes generally do not contain genes for T4P retraction ATPases^21^. Alternatively, retraction could also be conditional depending on experimental conditions.

### T4P promotes *H. volcanii* surface attachment in flow

Based on our visualization of steady tip contact, we suspected that T4P function in securing surface attachment. Consistent with this hypothesis, *H. volcanii* lacking all pilin isoforms fails to form microcolonies on plastic surfaces^25^. Mutants expressing a combination of these isoforms showed quantitatively distinct abilities to form these microcolonies, but how this is directly related to single cell adhesion is confounded by many factors of microcolony growth. To directly test the contributions of each pilin isoform in adhesion of single cell to surfaces, we used a flow-based microfluidic assay to directly measure their attachment strength. When attached to a surface, single cells in flow experience shear forces that tend to detach them from the surface^28^. Shear force increases linearly with flow velocity, which we here tune via flow rate, controlled by a syringe pump. In these microfluidic experiments with a channel width of 500 μm and height of 90 μm a flow rate of 100 μl/min exerts a force of approximately 10 pN on a 4 μm^2^ single cell at the channels centerline^29^. We injected an exponentially-growing culture in a microchannel, allowed cells to settle and attach to the glass surface, after which we initiated flow, increasing flow rate in a stepwise fashion (**Fig. 2A**). During this flow ramp, we continuously imaged surface attached population at the single cell resolution to count the number of cells remaining on the surface as a function of flow intensity.

**Figure 2:**
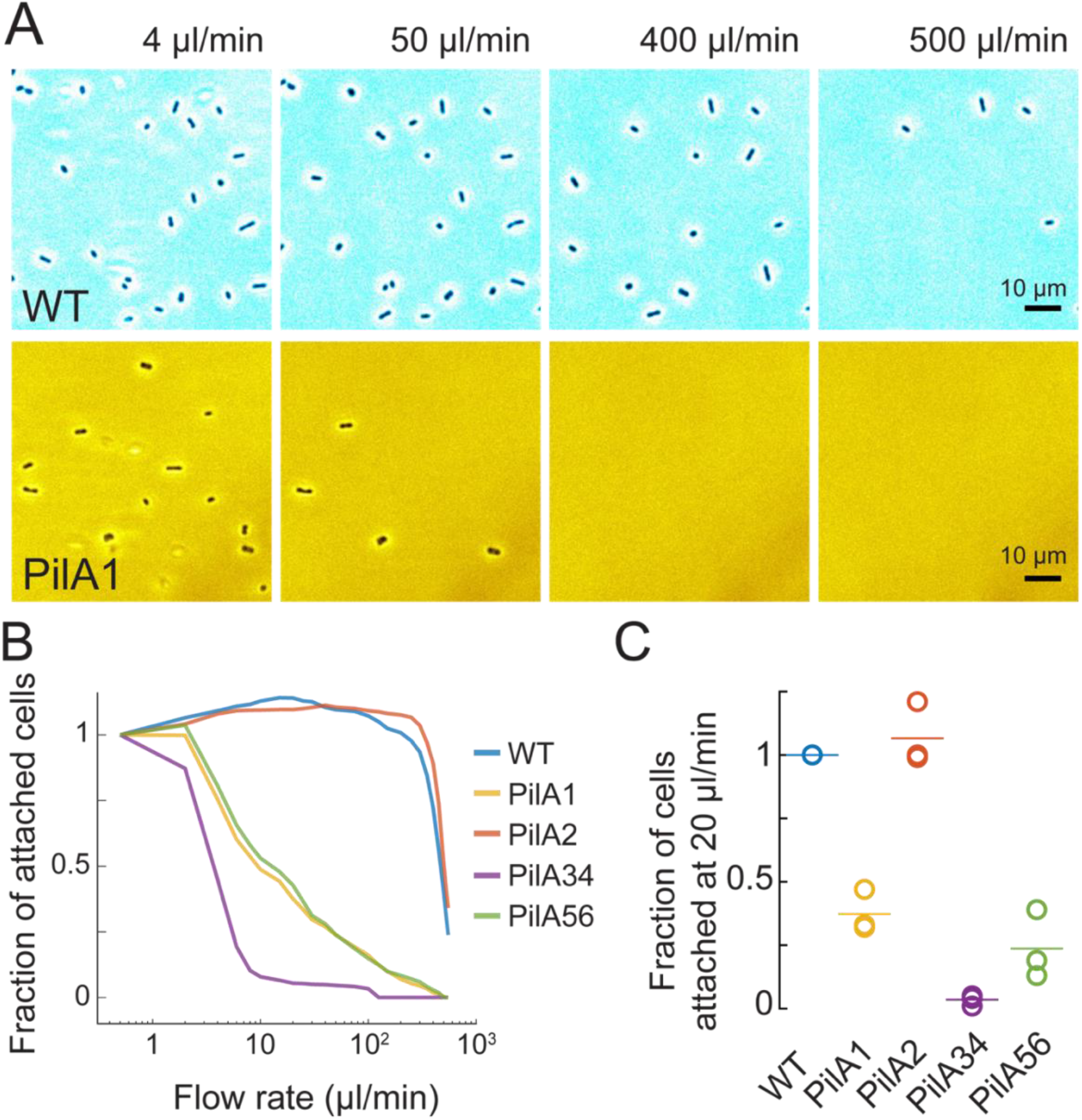
*H. volcanii* T4P enhances surface attachment. We incubated *H. volcanii* cells in microchannels to allow for attachment. We then incrementally increased flow rate in the channel every two minutes. **(A)** Phase contrast snapshots of WT and PilA2 strains at increasing flow rates. Most PilA2 cells gave already detached at 50 μl/min. (**B)** Fraction of surface-attached cells for strains expressing each pilin isoforms during an increasing ramp in flow rate. WT and PilA2 populations remain attached as shear stress increases, while cells from strains expressing the other PilA isoforms detach at lower flow rates. **(C)** Fraction of cells remaining attached at 20 μl/min. All fractions are normalized to the respective WT fraction of 3 independent experiments.

We found qualitative differences between WT and *pilA* mutants. WT cells remained attached to the surface at flow rates below 200 μl/min (∼ 20 pN shear force). PilA2 cells could resist flow forces as well as WT (**Fig. 2B**). By contrast, most pilin mutants detached at low flow rates (**Fig. 2AB**). The population of surface-associated cells in the PilA1, PilA3/4 and PilA5/6 strains dramatically decreased at flow rates near 20 μl/min corresponding to a maximum of ∼ 2 pN shear force, indicating a very weak adhesion (**Fig. 2C**). In summary, iSCAT imaging along with biophysical characterization of adhesion strength suggests that *H. volcanii* T4P play an important mechanical role in surface attachment. The characterization of mutants expressing single pilin isoforms shows that PilA2 is necessary and sufficient to phenocopy WT-level piliation and surface adhesion. The other pilin subunits are however not necessary for surface attachment. In addition, the low absolute value of adhesion force of mutants with low piliation indicate that *H. volcanii* exclusively uses T4P to attach on surfaces and doesn’t produce other unspecific surface adhesins such as polysaccharides in its planktonic state. Given their adhesive function and their importance in biofilm formation in bacteria, we hypothesized that T4P could promote *H. volcanii* biofilm formation on surfaces under flow. We thus first investigated the ability of WT *H. volcanii* to form immersed biofilms under flow conditions on long timescales.

### *H. volcanii* uses T4P to form clonal biofilms in flow

Biofilms form in flow when surface-attached single cells proliferate while maintaining cohesion with their progeny. To achieve this, they rely on cell-surface and cell-cell interactions. Given the large decrease in adhesion strength of mutants exhibiting low piliation, we tested whether T4P-dependent adhesion is crucial in the initiation of *H. volcanii* biofilm formation. We therefore first assessed WT *H. volcanii*’s ability to form biofilms in microfluidic channels maintained at 45 °C. We applied a shear stress of ∼ 0.5 pN which could still allow a few cells to attach in all isoform mutants (flow rate of 20 μl/min, corresponding to 5 μl/min in adhesion experiments). We found that single WT cells remained attached to the surface, grew and divided to form large multicellular structures (**Fig. 3A, supplementary movie 2**). In flow, microcolonies developed into mature biofilms nearly 100 μm-wide over the course of hours. Single cells eroded from the biofilm, thereby stretching the microcolonies in the flow direction to generate a “comet” shape (Figure 3B). This indicates that biofilm-dwelling archaea are loosely attached to the community after division. Ultimately, when biofilms reach a critical size, colonies ultimately slowly shear-off the surface to slowly roll down with the flow after 30 h of growth (**Supplementary movie 2)**.

**Figure 3:**
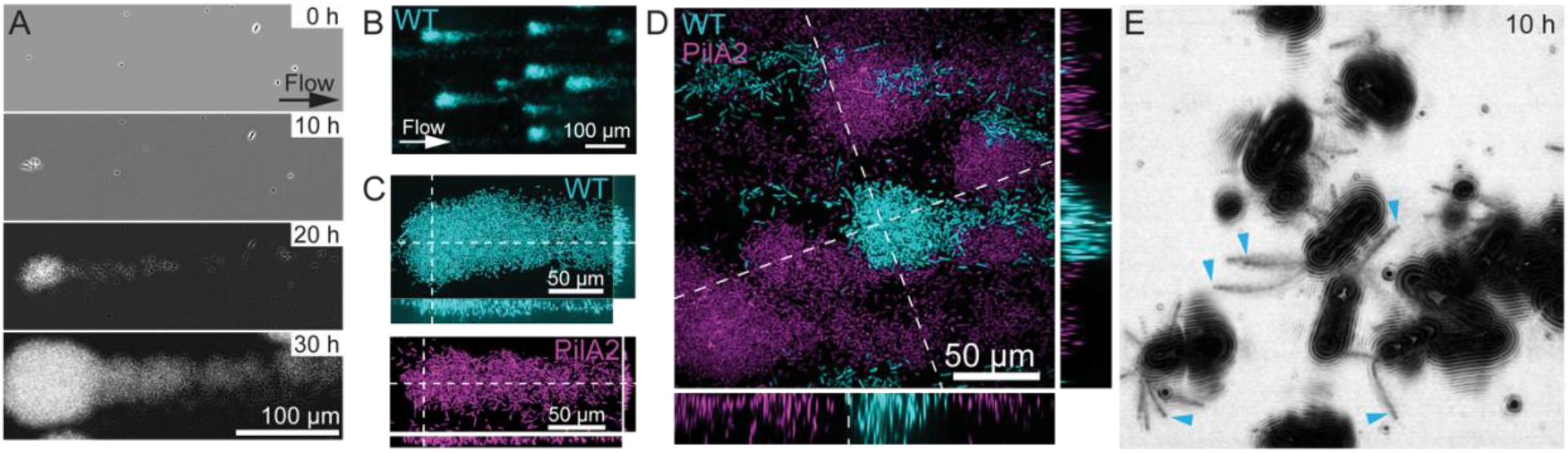
*H. volcanii* uses T4P to form biofilms in flow. **A)** Time series of phase images from WT biofilm growth under constant flow of 20 μl/min. *H. volcanii* forms comet-shape biofilms stretching in the flow direction. **B)** Confocal microscopy of WT-mCherry biofilms. **(C)** Confocal images of biofilms from WT-mCherry (top) and PilA2-GFP (bottom). Orthogonal views show that biofilms from both strains extend in the vertical direction. **(D)** Confocal visualizations of WT-mCherry and PilA2-GFP mixed biofilms. **E)** Representative iSCAT image of a microcolony early on during biofilm formation. The image corresponds to the standard deviation of 500 iSCAT images to improve visualizations. Blue arrowheads indicate T4P.

We inspected the three-dimensional architecture of mature biofilms of strains constitutively expressing fluorescent proteins by confocal spinning disk microscopy (**Fig 3B-D**). These visualizations allowed us to quantitatively compare the growth of strains expressing T4P subunit isoforms with WT. Out of the four mutants, only PilA2 consistently produced hundreds of biofilms as WT did (**Fig. S1**). In 2 out of 7 replicates, PilA1 formed a single biofilm, in 2 out of 6 attempts we found a single biofilm of the PilA3/4 mutant. PilA5/6 never produced a single biofilm, however a few cells managed to colonize the surface. Overall, these experiments revealed that strains with strong defects in T4P assembly are unable to form mature biofilms, while WT and PilA2 strains that are capable of forming T4P robustly formed multicellular structures. Since at this flow all mutants retain some ability to attach to the surface, our results are consistent with T4P mediating cell-cell attachment that is so critical to maintain biofilm architecture.

We asked whether PilA2 was sufficient to maintain *H. volcanii* biofilm architecture. We found that when grown separately, biofilms of WT and PilA2 strains showed seemingly identical architecture under flow (**Fig. 3C**). We however wondered whether lacking all other isoforms could create subtle changes that would impact architecture and fitness of the PilA2 isoform. To inspect potential differences, we performed competition experiments between WT and PilA2 strains each expressing a distinct fluorescent protein (**Fig. 3D**). Both biofilms extended in the vertical direction and again, there was no obvious difference in their architecture. In addition, we could not measure any growth advantage for WT over PilA2.

### *H. volcanii* physiologically adapts to the biofilm lifestyle

To better understand *H. volcanii* physiological changes associated with the multicellular lifestyle, we measured the transcriptional changes induced by biofilms growth. In particular, we wondered whether biofilm-dwelling archaea tend to increase the production of matrix components such as T4P to promote biofilm resilience. We quantitatively compared the transcriptomes of biofilm and exponentially-growing planktonic WT H26 populations. RNAseq data shows that most *pilA* isoforms tend to be upregulated in biofilms (**Fig. S2**). In agreement with adapting a sessile lifestyle, biofilm-derived cells exhibited a decreased expression of archaellum genes *arlA1, arlA2* as well as *arlD2* and *arlH* (**Table S1**). Consistent with these results, iSCAT visualization of nascent biofilms show that single cells extended many T4P early on as they nucleated biofilms (**Fig 3E**). Unfortunately, due to cell-crowding effects, we could not rigorously quantify T4P abundance at this stage, nor could we visualize them at later stages of biofilm formation. Transcriptomic and iSCAT results further validate a model where T4P play a crucial role in biofilm architecture and area actively produced to adapt to the multicellular lifestyle. 5 out of 6 pilins were upregulated in biofilms, indicating they play alternative functions within biofilms, or that these functions are condition-dependent.

### Stopping flow induces rapid biofilm dispersal

Our results show that T4P promote cell-surface adhesion, but how cell-cell cohesion is established in *H. volcanii* biofilms is still unclear. While the components and role of extracellular polymeric substances (EPS) in biofilm structural integrity has been identified for various bacterial species, whether a homologous matrix is formed in *H. volcanii* biofilms is less clear. Visualizations show the potential presence of polysaccharides and DNA in *H. volcanii* biofilms but how these contribute to cell-cell cohesion is not established^14^. The erosion of cells from the biofilm in the direction of flow suggests a rather loose connection between cells (**Fig. 3AB**). In addition, dynamic imaging of mature biofilms at high frame rate shows that while single cells are secured in the biofilm structure, their positions undergo slight fluctuations (**Fig. 4A, Supplementary movie 3**). This supports that biofilms are loosely cohesive via T4P and do not produce a polysaccharide matrix.

**Figure 4:**
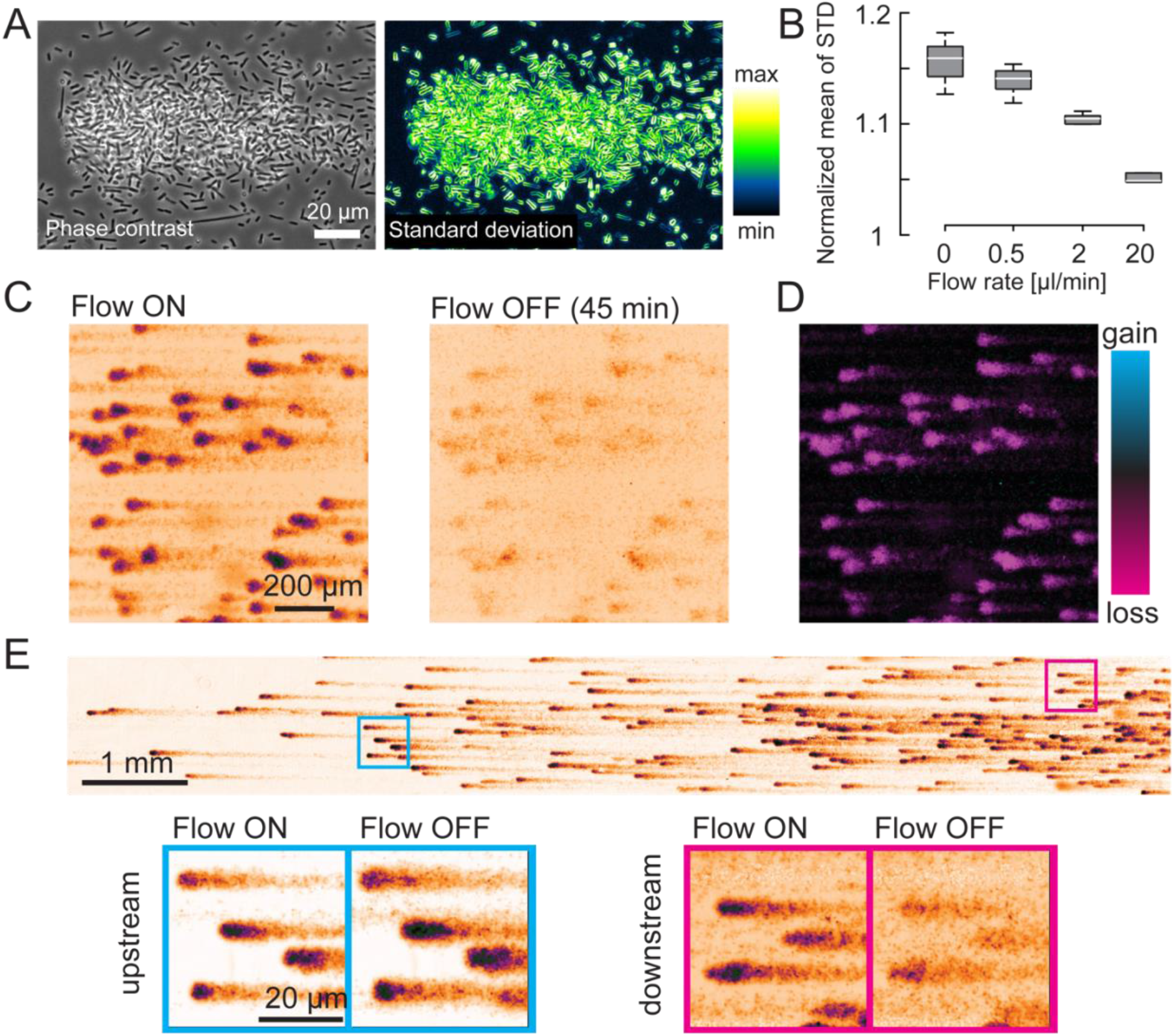
Flow promotes cell-cell cohesion in *H. volcanii* biofilms. **(A)** Phase contrast image of an H26 biofilm and standard deviation of phase-contrast movie of the same biofilm of biofilm shows that cells are stationary but wiggle in the community (**Supplementary movie 3**). **(B)** Mean standard deviation of biofilms phase contrast image sequences. The reduction of the standard deviation shows a stabilizing effect of flow. **(C)** Stopping flow disperses biofilms. Fluorescence images of WT-mCherry biofilms grown overnight at constant flow (20 μl/min). Flow was then stopped for 45 min and restarted for 5 min to flush out unattached cells before imaging. Biofilms largely dispersed during this procedure and few cells remained attached to the surface. **(D)** Relative fluorescence intensity changes upon flow arrest. **(E)** Biofilm dispersion is heterogeneous along the channel. Downstream biofilms disperse upon flow arrest, while upstream biofilms do not.

As a result of the apparently loose cell-cell interactions, we anticipated that biofilms would further erode with increasing flow. We however found that flow reduced the amount of fluctuations in the position of single cells (**Fig. 4B**). Thus, we hypothesize that hydrodynamic forces may stabilize cohesion between cells within biofilms. In these experiments, we found that that biofilms started dispersing when flow stopped. To rigorously characterize this effect, we grew biofilms overnight with an RFP expressing strain under flow. We then temporarily stopped flow for 45 min before imaging. Indeed, we found that the total fluorescent intensity dramatically decreased and only a fraction of the cells remained attached to the surface (**Fig. 4CE**). This suggests that flow is a crucial factor promoting structural integrity of biofilms.

Biomass loss was heterogeneous along the microchannel. Most biofilm dispersal occurred near the outlet, while biofilms localized upstream kept growing when flow was stopped. We compared the biofilm structural integrity between upstream and downstream sections. We visualized 20 h-old biofilms while decreasing flow rate in a stepwise manner between 20 μl/min and 0 μl/min (**Fig. 4E**). We found that while biofilms in the upstream section grew at all flows including at 0 μl/min, downstream biofilms dramatically dispersed only when flow rate was set to zero (**Fig. 4E**). High resolution visualizations of the dispersed biofilm show that the colony lost a large fraction of cells but the biofilm base left a cell imprint at the coverslip surface, indicating cell-surface interaction is maintained to some extent.

We then wondered what distinguishes upstream and downstream population so that only colonies near the channel outlet disperse. In immersed biofilm experiments under flow, cell density commonly increases along the channel, as flow removes single cells from upstream biofilms that can reattach and grow downstream (**Fig. 4E**)^30^. We hypothesized that local cell density might influence the dispersal process. Consistent with this, we found that areas with strong dispersion were also more colonized. To verify this dependence on density, we observed dispersal of biofilms grown to different densities where colonies are distinctively smaller. High density biofilms dispersed in the downstream section when flow was stopped. By contrast, biofilms of lower density did not disperse upon flow arrest, but rather uniformly increased in biomass (**Fig. S3A**). In summary, these results suggest that flow has a direct effect on biofilm structural integrity. In the absence of flow, a destabilization of cell-cell interactions loosens biofilm cohesion. This loss of attachment liberates archaeal cells in the fluid which may further stimulate dispersion enabling interactions with planktonic cells.

We envision a mechanism where T4P maintain connections between cells under the influence of flow-forces. These interactions become weaker when flow stops allowing cells to separate from each other, explaining the density-dependent phenotype. Based on this rationale we hypothesized that exogenously introducing cells within the medium by perfusing the microfluidic channel would be sufficient to promote dispersal of biofilms that would normally not disperse upon flow arrest (**Fig. S3B**). We thus stopped flow of grown biofilms, injected an exponentially growing cell culture in fresh medium in the microchannel and compared dispersion with a negative control (medium only). Incubation of WT biofilms with liquid grown WT cells vastly increased dispersion compared to the medium control homogenously along the entire channel (**Fig. S3CD**). To test the extent to which T4P of newly-introduced cells interfered with T4P providing biofilm cohesion, we performed a similar experiment using a T4P-lacking mutant. We incubated WT biofilms with an exponentially growing culture of a deletion mutant lacking the prepilin peptidase gene *pilD*, which in turn prohibits pilin subunit polymerization. We still observed that planktonic *ΔpilD* cells promoted dispersal over a blank medium control, albeit to a lower extent compared to WT cells (**Fig. S3D**). Therefore, planktonic cells can perturb the cohesion between biofilm-dwelling cells, and T4P may improve this disruption possibly by further disengaging bound T4P from their biofilm targets.

## Discussion

Biofilms represent the most common lifestyle for many unicellular organisms. Surface appendages play a dominant role in surface attachment, biofilm formation and integrity. T4P facilitate biofilm morphogenesis in *P. aeruginosa, V. cholerae* and *N. meningitidis*. Here, using high resolution label-free microscopy imaging of T4P, in conjunction with biofilm growth experiments under controlled flow conditions using a microfluidic setup, we find that flow-dependent T4P-mediated surface adhesion and pili-pili interactions are essential components for biofilm formation in *H. volcanii*.

At the level of T4P biogenesis, we found that PilA2 alone is sufficient to phenocopy WT piliation and surface adhesion properties. This suggests that the other pilin isoforms do not play a fundamental role in initial surface adhesion under flow. Interestingly, PilA5/6 were upregulated in matured biofilms in our microfluidic setup under flow. This is in agreement with the study from Esquivel *et al*.^25^ in which different *H. volcanii* mutants were tested for their adherence to a glass surface under static (non-shaking) conditions. Under these conditions, PilA2 was not important for microcolony formation, but PilA5/6 mutants produced the most microcolonies. Therefore, we postulate that these pilin play different roles for biofilm formation during different environmental conditions: PilA2 is the main adhesive pilin under flow conditions, whereas PilA5/6 are important without flow or at later stages of biofilm morphogenesis.

We showed that *H. volcanii*’s can form biofilms in a constant flow environment. Our results highlight the functional role T4P for *H. volcanii* biofilm formation, by both acting on cell-surface adhesion and cell-cell cohesion. Inspired by the observation that cells didn’t seem to be lock into a position, we postulate that T4P loosely maintain *H. volcanii* biofilm structural integrity. This scenario is reminiscent of *N. meningitis* biofilms where pili-pili interactions promote cell-cell cohesion that maintains biofilm integrity^31^. In this case, T4P retraction strengthen these interactions, but *H. volcanii* T4P do not appear to retract. However, flow can induce tension force in T4P, thereby stimulating these interactions, possibly in a mechanism that involves catch bonds between T4P. Our results suggest that flow maintains biofilm structural integrity is consistent with this mechanism. In addition, stopping flow stimulates dispersal, most likely in a passive mechanism where lack of tension forces in T4P loosen their connections. Flow-dependent dispersal indicates that for *H. volcanii*, the biofilm lifestyle confers a selective advantage in flow but not in a resting fluid. When flow stops, cohesion between cells is irreversibly lost, and disassociated cells move away from the biofilm. Flow may thus represent a mechanical signal, indicating that conditions are favourable to adopt a planktonic lifestyle and explore other environments. Conversely, flow may indicate that there exists a locally favourable flux of nutrients, for which maintaining a sedentary lifestyle is more efficient.

Taken together, our study identifies T4P and flow as key components in the process of biofilm development in *H. volcanii*. Improving visualization methods within biofilms will provide more information on their structural role in thee multicellular communities. More generally, we envision that replicating different facets of the natural environments of archaea win the lab will enable of more comprehensive understanding of their physiology.

## METHODS

### Strains and culture conditions

The *Escherichia coli* cells used for cloning were grown in LB-medium^32^ supplemented with 100 μg/ml ampicillin under constant shaking at 150 rpm at 37°C or LB-agar plates containing ampicillin at 37°C. *Haloferax volcanii* cells were either grown in full YPC-medium^33^ when used for transformation or selective Ca medium^33^ supplemented with an extended trace element solution ^34^ for experiments. *H. volcanii* plates with solid medium for transformants were prepared as described by Allers *et al*.^33^. *H. volcanii* was grown in a shaking incubator at 45°C, plates were sealed in plastic bags to prevent evaporation and were grown at 45°C as well. In general, a pre-culture of relevant *H. volcanii* strains were grown in 3-5 ml Cab medium in a rotational shaker at 45°C overnight and where required, diluted the next day to reach the required density at the time of experiments. Strains used for this study are indicated in Table S1

### Plasmid construction

DNA modification enzymes and PHUSION® polymerase for DNA amplification were all obtained from New England Biolabs (NEB) and were used as indicated by the manufacturer. Knock-out plasmids for the deletion of the different pilins and the major archaellin ArlA1 were cloned by the amplification of a ∼500 bp upstream region and ∼500 bp downstream region of the respective genes. For the pilin knock-out plasmids the amplified upstream regions of the different pilins contained a KpnI restriction side and at the 3’-end a region homologous to the respective downstream fragment. The downstream fragments were amplified with a XbaI restriction site. The different fragments were digested with either KpnI or XbaI and subsequently cloned into integrative plasmid pTA131 opened with KpnI and XbaI as well. The ligation products were transformed into *E. coli* Top10 for circularization. The up- and downstream fragment for the arlA1 knock-out plasmid were connected by a NdeI restriction side and were cloned in pTA131 via KpnI and XbaI. Plasmids and primers used in this study are indicated in Table S1.

### Plasmid transformation and strain construction of *H. volcanii*

Before transformation of the constructed knock-out plasmids into *H. volcanii* they were passed through a dam/dcm deficient *E. coli* strain. *H. volcanii* transformation was performed using polyethylene glycol 600 as described before^35^. Transformants were plated on selective Ca-plates and the plates incubated at 45°C until colonies were visible. To generate the different knock-outs one colony of per knock-out attempt was transferred to 5 ml YPC-medium. When the cultures reached OD_600_ 1 they were diluted back 1:500 into 5 ml fresh YPC-medium. This step was repeated in total three times. To screen for pop-out events the single cultures were diluted hundred times and 100 μl of each dilution plated on Ca-plates supplemented with 50 μg/ml 5-fluoorotic acid and 10 μg/ml Uracil. Colonies from these plates were screened via colony PCR for gene deletions with the primers indicated in Table X. To obtain the strains with just one set of pilin the others were deleted consecutively. Subsequently to remove the archaellum, the major archaellin ArlA1 was deleted in all pilin deletion mutants following the process described above.

### Media preparation

Stock solutions were prepared as follows. For 2 L of a 30% buffered Salt Water (BSW) stock solution 480 g NaCl, 60 g MgCl_2_·6 H_2_O, 70 g MgSO_4_·7 H_2_O, 14 g KCl, 40 mL were first dissolved in ∼1.6 L warm distilled water, then 40 mL of 1M Tris-HCl pH 7.5 was added and topped up with distilled water to a final volume of 2 L. 10x CA stock solution was prepared by dissolving 10 g of Casamino Acids to ∼150 mL of distilled water and stored at 4 °C. Then the pH was adjusted to 7.2 by addition of 1 M KOH. Distilled water was then added to a final volume of 200 mL. 1000x vitamin solution was prepared by dissolving 50 mg Thiamine and 5 mg Biotin in 50 mL of distilled water. 100x trace element solution was prepared by (need details here).

To prepare medium, 200 mL of BSW, 100 mL of distilled water, 33 mL of 10x CA stock and 2 mL of a 0.5 M CaCl_2_ were mixed. This solution was then autoclaved and/or filtered using 0.22 μm medium filter units. Immediately before using media vitamins and trace elements were added at appropriate dilutions, 1000x and 100x, respectively.

### Microfluidic device

Microfluidic devices were prepared by baking a PDMS and curing agent mixture (Sylgard 184, Dow Corning) at a ratio of 10:1 at 80°C for at least 6 hours using microfabricated one dimensional channels arranged on a silicon wafer^36^. Individual devices were cut out and liquid reservoir with diameter 6-8 mm at inlet, and in- and outlet for tubing connections were punched (see schematic Figure 4A). Devices were then bonded onto coverslip glass slide which were previously cleaned by rinsing with isopropanol in between two water-ethanol-water rinsing steps. Drops of water was blown away with compressed air followed by a short incubation at 80°C. To bond coverslip glass and PDMS devices together, both surfaces were oxygen plasma treated (Zepto, Diener Electronic) for 1 min beforehand. A round glass coverslip was then bonded using the same plasma surface treatment ontop of the PDMS devices to cover the punched out liquid reservoir which functioned as a bubble trap. Devices were then incubated at 80°C briefly and immediately used for experiments. For biofilm growth experiments microfluidic channels with a width of 2 mm and height of 300 μm were used. For iSCAT and surface adhesion assessments smaller channels without a liquid reservoir with a width of 500 μm and height of 90 μm, respectively, were used.

### iSCAT

Cells from liquid pre-culture were diluted to reach ∼OD 0.1 the next morning. Cells were then diluted (∼3-6x) and used for iSCAT imaging between OD 0.15-0.2. Cells were added to small microfluidic channel at low flow using kd science syringe pump. The observation chamber was heated to the maximum temperature (∼42 °C). iSCAT experiments were performed as described in detail previously^27^. Typically, iSCAT movies were recorded for a duration of 10 s at 50 fps. Quantification of surface appendages from post-processed iSCAT movies were analysed by randomly displaying an iSCAT movie in Matlab without knowledge of the strain. Pili were then identified visually for each cell.

### Adhesion

A pre-culture of cells from a plate was grown overnight. The next day, the culture was diluted accordingly to reach an OD between 0.15-0.2 for adhesion experiments. A microfluidic channel (width: 500 μm; height: 90 μm) was perfused with medium, then 100 μl of a cell solution was added to the luer stub (27G x ½” Luer Stubs, Instech) connecting the syringe outlet nozzle and tubing. The liquid level was allowed to drop slightly by gravitational force before a small volume of fresh medium was added on top and the luer stub was reconnected to the syringe making sure no air bubble got introduced. Flow was set to 20 μl/min for 1-2 min till cells appeared in the microfluidic channel, then flow was stopped to allow cells to adhere to the glass for maximal 15 min. Then, flow was restarted at 0.5 μl/min, and once it appeared steady acquisition was started. A phase image was acquired at one location every 10 s using a 40x Ph2 objective on a Nikon Eclipse Ti inverted optical microscope equipped with a Prime95B sCMOS camera (Photometrics). The flow rate was increased every 2 min as indicated in Figure 2A until no more cells remained attached in the field of view. Then the procedure was repeated with subsequent strains at the same location. Image analysis was done in ImageJ. For the background subtraction, the last empty image in the sequence was used. First, a positive offset was applied to the entire image dataset, then the original last image was subtracted from the entire dataset providing a flat background intensity across the whole dataset. Number of cells in each image were then counted by thresholding background corrected phase contrast images and counting the particles using ImageJ. We computed shear stress from flow rate as in reference 29.

### Biofilm growth

For biofilm growth experiments tubing (1.09 mm outer diameter polyethylene tube, Instech) connected to a 30 or 50 ml plastic syringe (BD Plastipak) filled with Cab medium which was prewarmed in the microscopy heating chamber before loading the syringe. A blunt luer stub was used to connect syringe and tubing. The syringe pump (kd science) was connected to the inlet of a large channel microfluidic device. The liquid reservoir (with intentionally leaving a small air bubble in the reservoir) and observation channel were filled with medium before outlet tubing was connected and left to equilibrate for some time with flow. Then the inlet and outlet tubing were disconnected such that a small drop of medium was covering the outlet opening. Then 5-8 μl of a diluted or non-diluted pre-culture cell solution was pipetted into the microfluidic channel from the outlet filling up ∼75 % of the channel volume, making sure no cells reached the liquid reservoir at the inlet. The outlet was then closed with tape for 5-10 min to allow cells to attach to the glass before the inlet tubing was reinserted. After a short time (<3 min) the tape at the outlet was removed and the tubing reconnected. The cell coverage was then inspected by microscopy and excessive cells removed by repeatedly tapping the bottom of the microfluidic channel until the desired coverage was reached. Random images along the channel were recorded either before or just after cell loading, which were used for background subtraction for the analysis later. Microscopic observation was performed using a Nikon inverted microscope with temperature-controlled incubation chamber set to 45°C.

### Biofilm dispersal

1 mL of exponentially growing cells were centrifuged for 2 min at 16’100 rcf, supernatant was discarded and cells resuspended in an appropriate volume of fresh Cab medium to an OD_600_ of 0.2. A 1 mL BD Plastipak syringe with a blunt luer stub was filled with this cell solution, mounted on a syringe pump and used for biofilm dispersal experiments. To minimize flow disturbance during switching the inlet tubing from the large syringe used for growing biofilms overnight to the syringe containing the cell solution the inlet tubing was cut close to the luer stub and connected to the syringe containing the cell solution. The cell solution was pumped for 10 minutes making sure the entire channel was filled with incoming cells. Fluorescence images along the microfluidic channel were recorded with a 10X/0.25 Ph1 DL objective (Nikon) before and after biofilms have been exposed to flow manipulations as indicated. Generally, before recording the second image, flow was restarted for 5 min to flush out any non-attached cells.

### Fluorescence image analysis

Images were then analyzed as follows: background images of the empty channels acquired before biofilms were grown, were opened as a stack in ImageJ and the median projection was used as to flatten the background. The images were then transformed into a montage using ImageJ. A threshold was applied to the montage images in ImageJ to generate a mask of the cell area, and for each column of pixels (perpendicular to the direction of the channel) the median of the background was set to 0 by addition or subtraction in Matlab. Then, the overall median intensity of the background of the montage was set to 0 by addition or subtraction in ImageJ. This rendered an even flat background of median value 0 along the entire channel. The total fluorescence intensity was then obtained by masking the background area and summing the integrated intensity of the biofilms. The change in fluorescence intensity was calculated by subtracting the image acquired before the procedure from the image acquired after the procedure. The resulting differential image was then color coded to represent positive values in cyan and negative in magenta at equal intensity ranges. To calculate the fluorescence along the channel the intensity of the non-masked along the montage was summed for each column of pixels. The values were then smoothed by applying a moving average over 200-pixel columns.

### Measurements of cell fluctuations in biofilms

Biofilms were grown overnight as described above. Images (200 frames within 5 s) were acquired at 20 μl/min before flow rate was changed to 2, 0.5, 0 and back to 20 μl/min, respectively. The procedure was then repeated once more at the same location. Image sequences were acquired for each flow rate from which we computed standard deviation of image stacks using ImageJ. One sequence of flow ramps lasted about 4 minutes.

### RNA isolation

Biofilms grown for ∼20 hours were harvested as follows. Outlet tubing was disconnected and outlet opening was closed with tape. The inlet tubing was cut with scissors to minimize disturbances and the device was removed from the microscope incubation chamber. A second hole was punched from the sidewall into the liquid reservoir at the inlet and the reservoir liquid was carefully sucked out with a pipet, leaving only the microfluidic channel filled with medium. Then, the liquid reservoir was filled with 20 mM Hepes, 2% SDS at pH 8, the scotch tape at the outlet removed and liquid sucked through the channel from the outlet opening to detach the cells by gently pipetting a fraction of the Hepes solution back and forth. Finally the entire liquid was collected and transferred to an Eppendorf tube which was then vortexed to ensure cell lysis. Harvesting biofilm grown cells was completed within less than 5 min per channel. Once cells were harvested, an appropriate amount of Trizol and chloroform reagent was added to the Eppendorf tube according to the manufacturer’s protocol (protocol). After centrifugation at 4°C at 12’000 rcf for 15 min, the clear aqueous phase was transferred to a fresh nuclease free tube and RNA was isolated using Zymoresearch Nucleic acid concentrator column protocol according to the manufacturer’s protocol (details).

Cells form liquid culture were harvested from a diluted pre-culture cell suspension the following day at an OD_600_ of 0.13-0.15. Cells were centrifuged in 50 ml falcon tubes at maximum speed (xspeed) for 4 minutes and resuspended in a HEPES/SDS mix (as above) after discarding supernatant. Then the same protocol as for biofilm-isolated cells was followed.

RNA was then analyzed using bioanalyzer and rRNA depletion was done using siRNAtools depletion assay for *H. volcanii* based on magnetic removal of rRNA. The samples were then processed at genomics facility EPFL. Transcriptomics analysis was done using usegalaxy.org. Using hisat2 map reads to genome, paired-end library, reverse direction. Count the maps using htseq-count tool. We analyses differential expression using the deseq2 tool. Genome sequence <u>ASM2568v1 - Genome - Assembly - NCBI (nih.gov)</u>

## Supplementary Material

**Figure S1:**
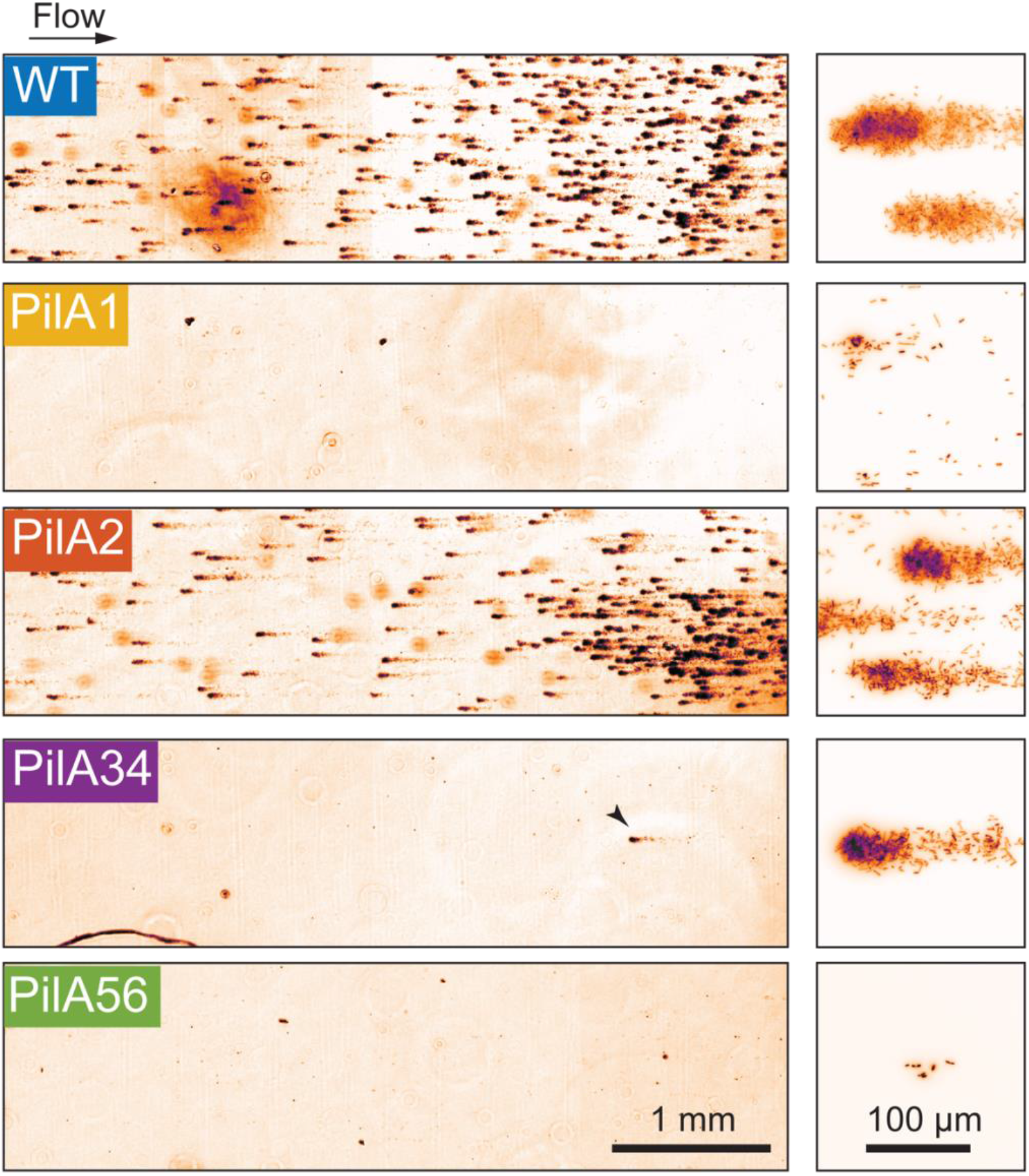
Representative fluorescence microscopy images of biofilms from strains expressing the respective PilA isoforms growing in flow. Only strains reliably producing T4P (WT and PilA2) can reproducibly form robust biofilms in flow. Biofilm formation is abolished in strains with low piliation (PilA1, PilA3/4 and PilA5/6). In these backgrounds, single cells occasionally attach to the surface, but rarely grow into mature biofilms. Right column: Close-up views of selected region in microfluidic channel.

**Figure S2:**
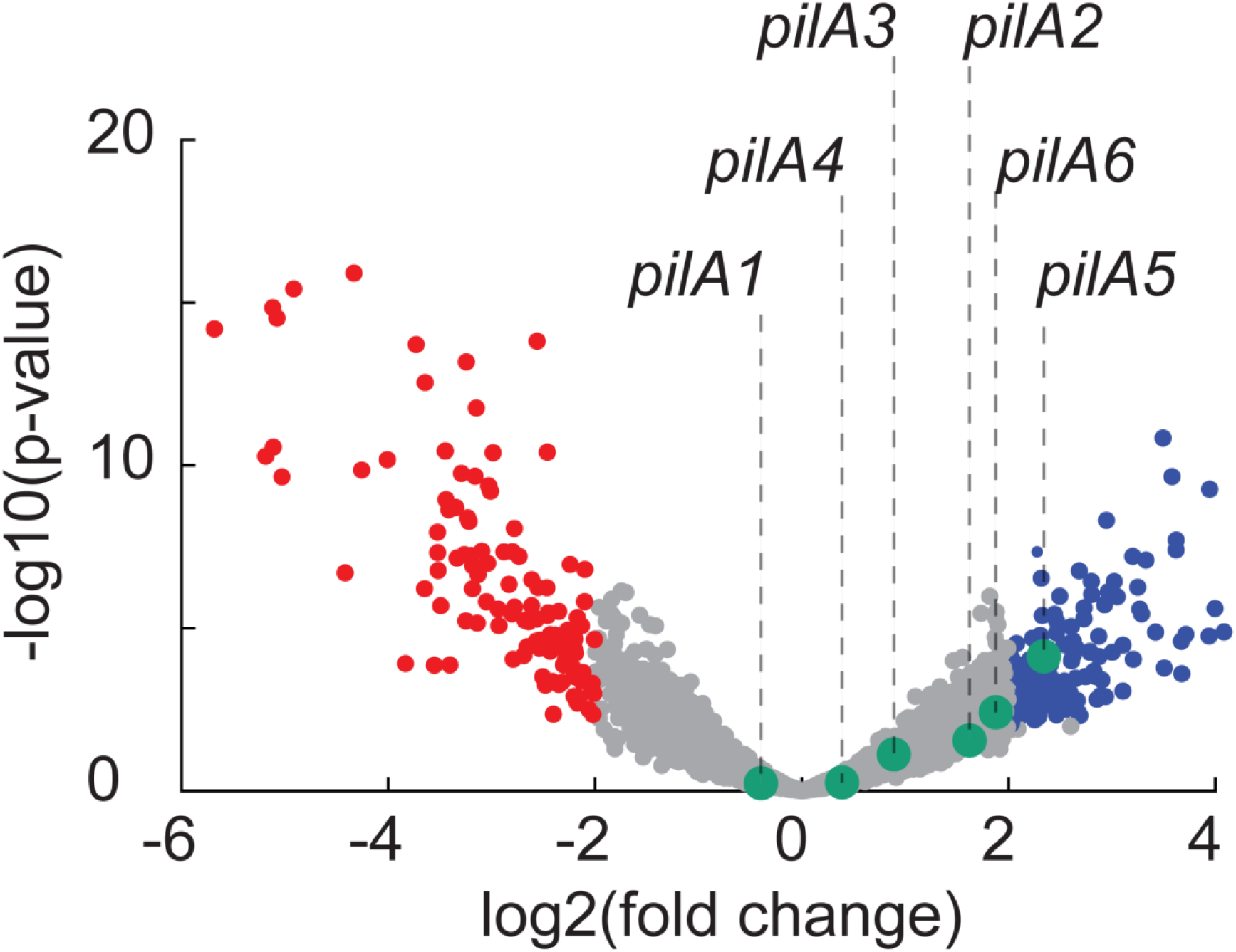
Volcano plot of gene expression measurements of WT biofilms relative to exponential liquid culture. *pilA5* is clearly upregulated in biofilms, while other *pilA* subunits tend to be upregulated, albeit with a low p-value. The relatively low values of fold-changes in gene expression in a well-documented phenomenon in archaea.

**Figure S3:**
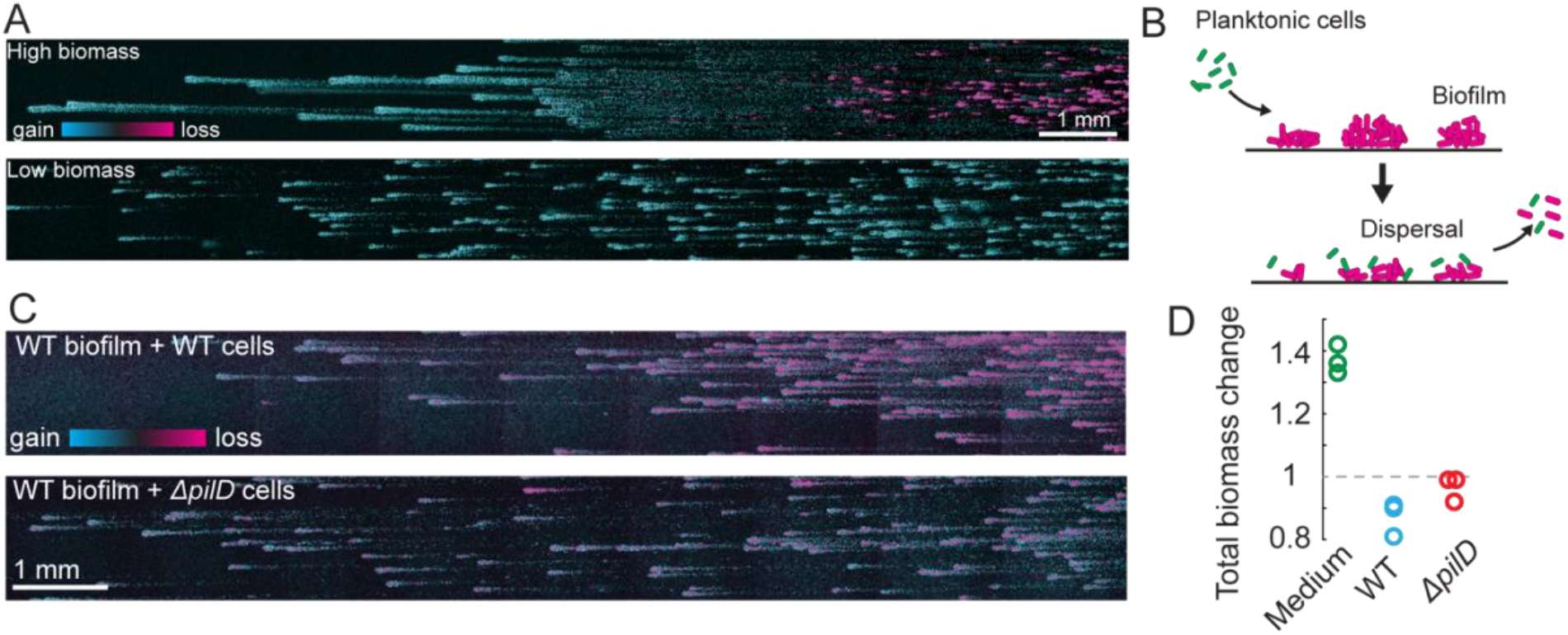
Interactions with planktonic cells stimulates biofilm dispersal. **(A)** Relative changes in biofilm biomass. Biofilms all grow in the upstream region while they disperse downstream. In conditions where biofilm biomass is lower, stopping flow does not induce dispersal. **(B)** To test the contribution of loose cells on dispersal, we introduce exponential-phase cultures in fresh medium into the channel containing mature biofilms and stop flow for 45 min. **(C)** Images showing change of fluorescence intensity upon incubation with liquid-grown cells. Liquid-grown WT cells were flushed into microfluidic channels containing WT biofilms. Unlike previous experiments shown in Fig. 4, biomass decreased throughout the entire microchannel. When incubated with the T4P-deficient *ΔpilD* mutant, biofilms also dispersed slightly. **(D)** Quantification of relative biomass change upon flow arrest.

**Table S1:** Strains used in this study

| Strain Name | Background strain | Genotype | Source/reference |
| --- | --- | --- | --- |
| ***H. volcanii*** |  |  |  |
| H26 | - | Δ*pyrE2* | 33 |
| HTQ 265 | H26 | Δ*pilA2*, 3-4, 5-6 (*pilA1* only) Δ*arlA1*<br>Δ*pyrE2* | This study |
| HTQ 266 | H26 | Δ*pilA1*, 3-4, 5-6 (*pilA2* only) Δ*arlA1*<br>Δ*pyrE2* | This study |
| HTQ 267 | H26 | Δ*pilA1*, 2, 5-6 (*pilA3-4* only) Δ*arlA1*<br>Δ*pyrE2* | This study |
| HTQ 268 | H26 | Δ*pilA1*, 2, 3-4 (*pilA5-6* only) Δ*arlA1*<br>Δ*pyrE2* | This study |
| ***E. coli*** |  |  |  |
| 10-beta Competent Cells "TOP10" | - | $\Delta$(*ara-leu*) 7697 *araD139 fhuA*<br>Δ*lacX74 galK16 galE15*<br>*e14- ϕ80dlacZΔM15 recA1 relA1*<br>*endA1 nupG rpsL (Str<sup>R</sup>) rph spoT1*<br>$\Delta$(*mrr-hsdRMS-mcrBC*) | New England Biolabs |
| *dam/dcm* Competent | - | *ara-14 leuB6 fhuA31 lacY1 tsx78*<br>*glnV44 galK2 galT22 mcrA dcm-6*<br>*hisG4 rfbD1 R(zgb210::Tn10) Tet<sup>S</sup>*<br>*endA1 rspL136 (Str<sup>R</sup>) dam13::Tn9*<br>(*Cam<sup>R</sup>*) *xylA-5 mtl-1 thi-1 mcrB1 hsdR2* | New England Biolabs |

**Table S2:**
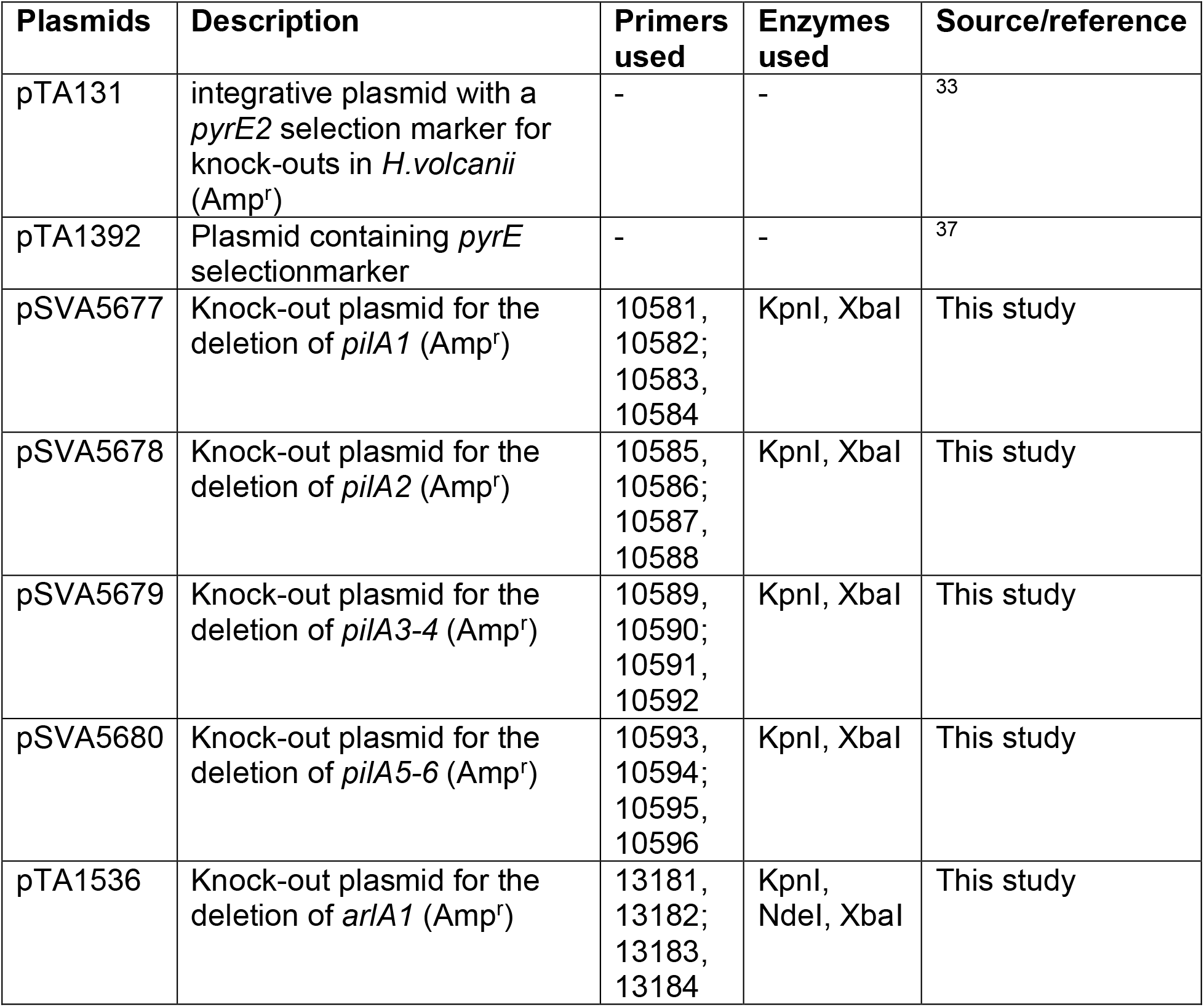
Plasmids used in this study

| Plasmids | Description | Primers used | Enzymes used | Source/reference |
| --- | --- | --- | --- | --- |
| pTA131 | integrative plasmid with a *pyrE2* selection marker for knock-outs in *H.volcanii* (Amp<sup>r</sup>) | - | - | 33 |
| pTA1392 | Plasmid containing *pyrE* selectionmarker | - | - | 37 |
| pSVA5677 | Knock-out plasmid for the deletion of *pilA1* (Amp<sup>r</sup>) | 10581,<br>10582;<br>10583,<br>10584 | KpnI, XbaI | This study |
| pSVA5678 | Knock-out plasmid for the deletion of *pilA2* (Amp<sup>r</sup>) | 10585,<br>10586;<br>10587,<br>10588 | KpnI, XbaI | This study |
| pSVA5679 | Knock-out plasmid for the deletion of *pilA3-4* (Amp<sup>r</sup>) | 10589,<br>10590;<br>10591,<br>10592 | KpnI, XbaI | This study |
| pSVA5680 | Knock-out plasmid for the deletion of *pilA5-6* (Amp<sup>r</sup>) | 10593,<br>10594;<br>10595,<br>10596 | KpnI, XbaI | This study |
| pTA1536 | Knock-out plasmid for the deletion of *arlA1* (Amp<sup>r</sup>) | 13181,<br>13182;<br>13183,<br>13184 | KpnI,<br>NdeI, XbaI | This study |

**Table S3.**
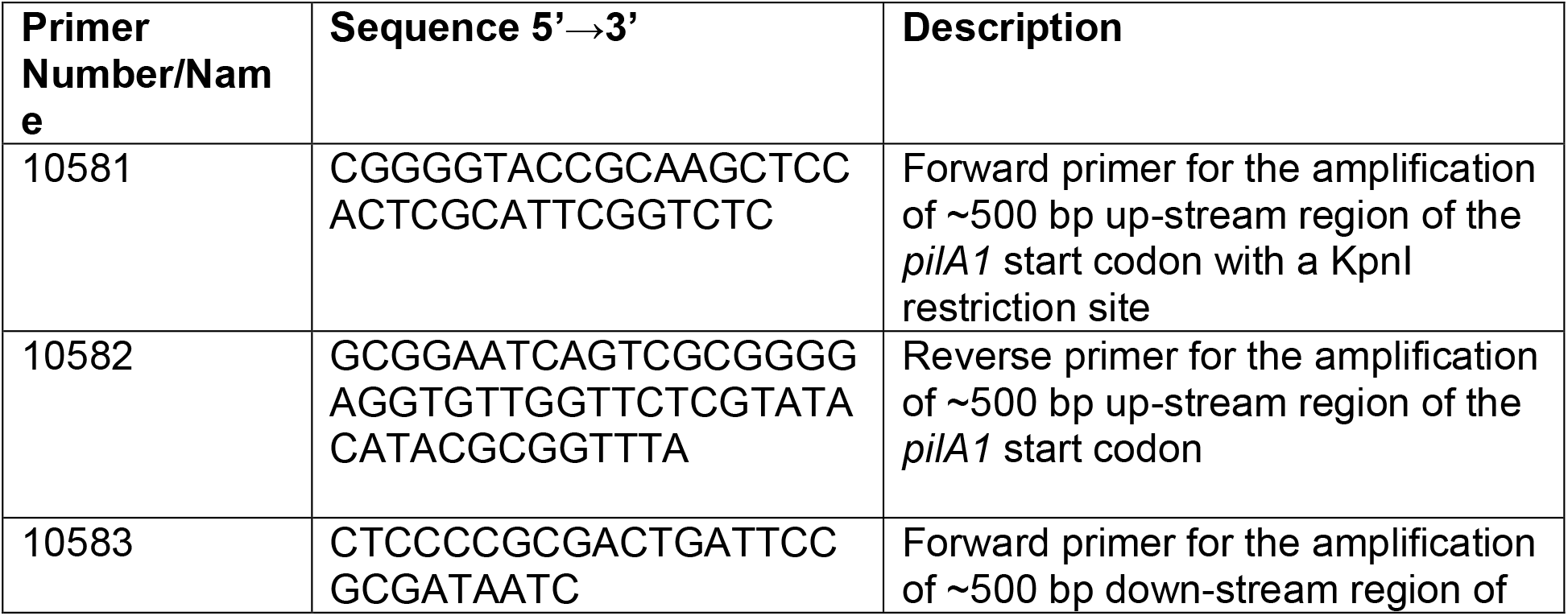
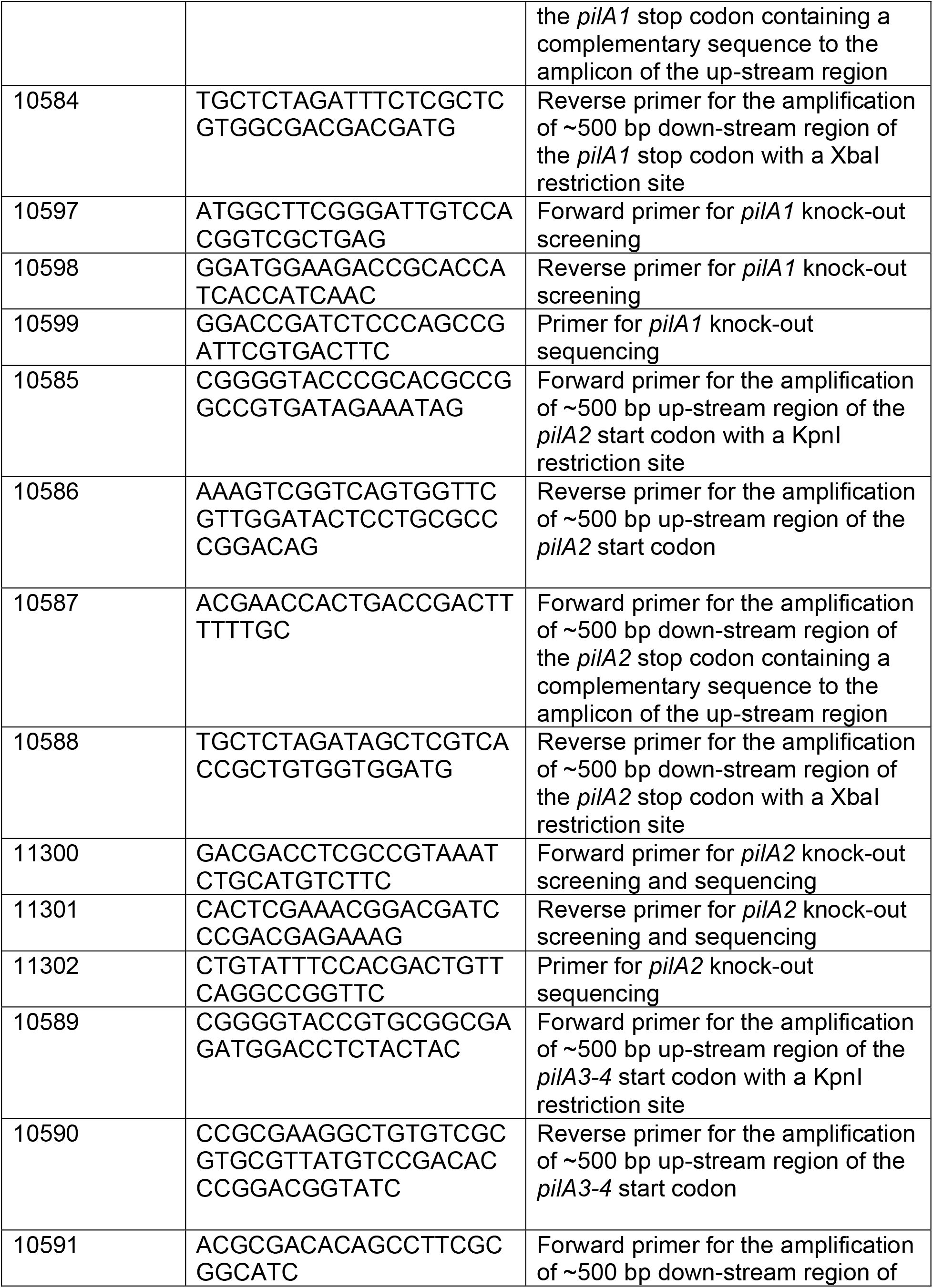
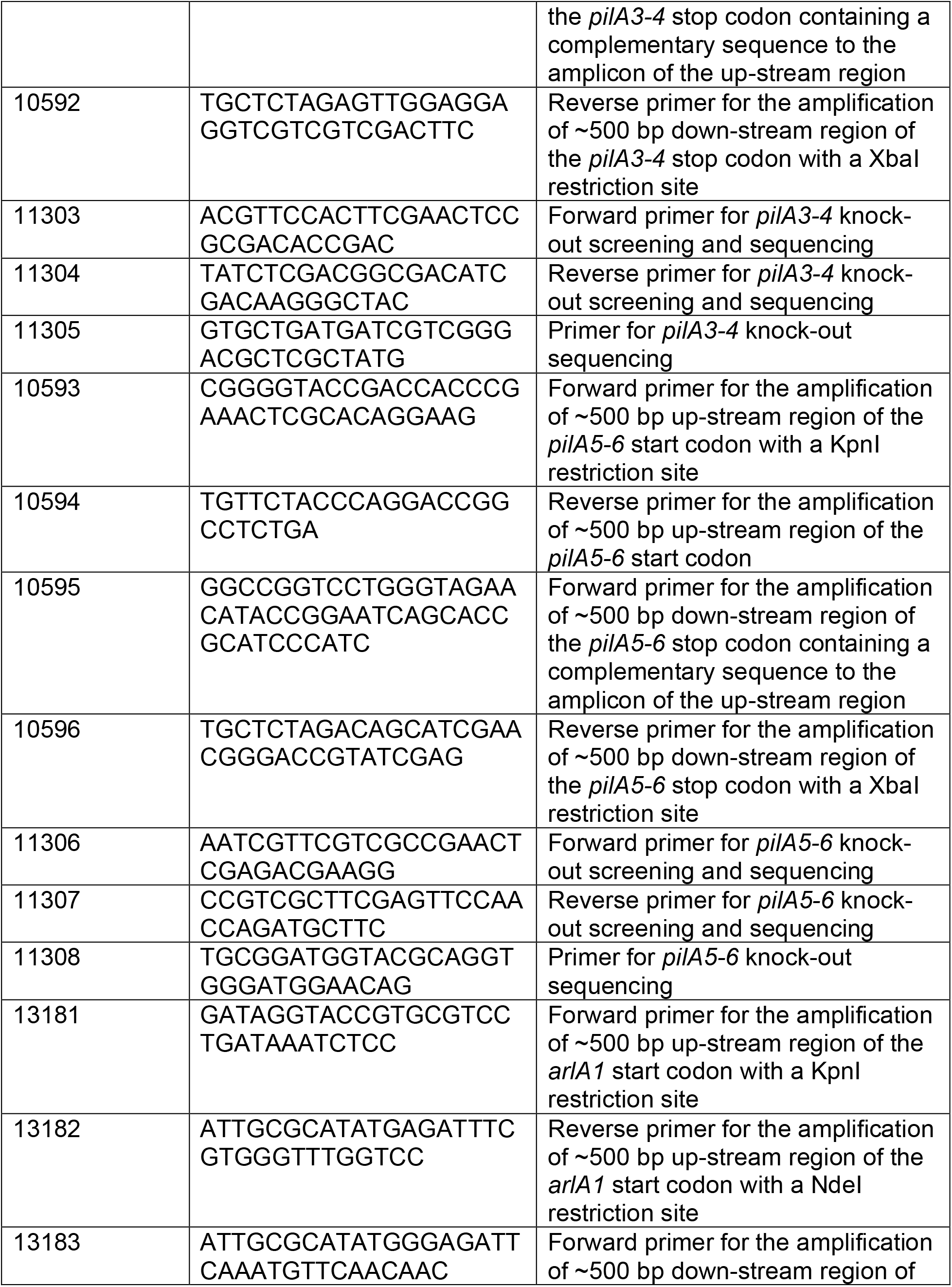
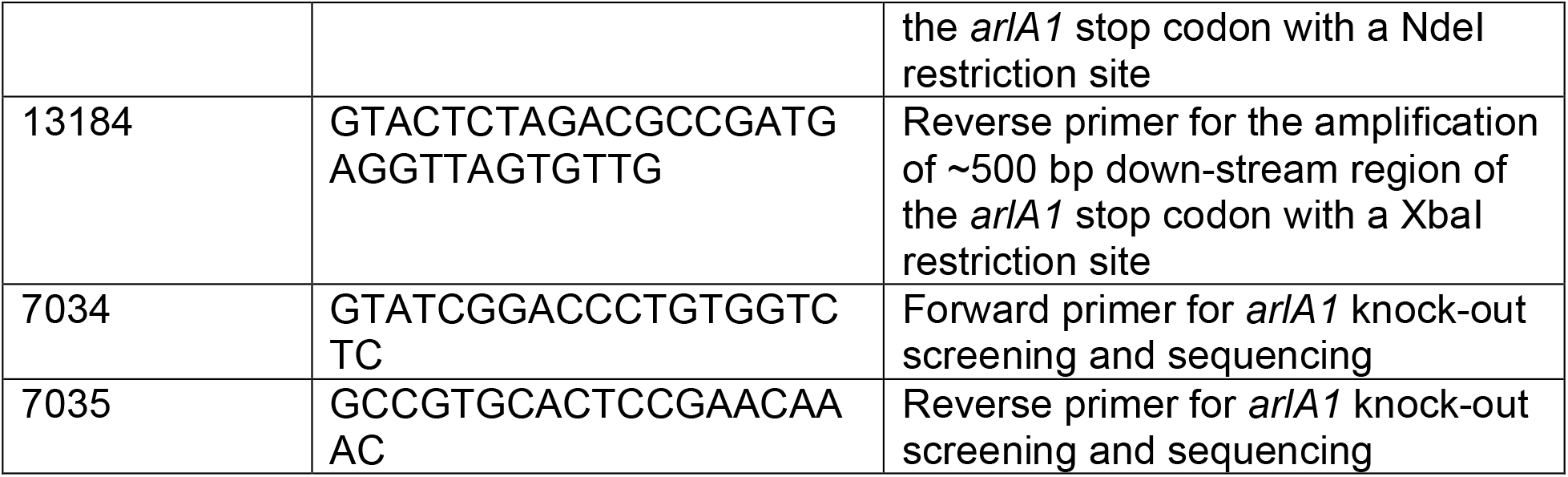
Primers used in this study.

## Notes

### Competing Interest Statement

The authors have declared no competing interest.

